# IL-15 synergizes with CD40 agonist antibodies to induce durable immunity against bladder cancer

**DOI:** 10.1101/2023.01.30.526266

**Authors:** Jeffrey L. Wong, Patrick Smith, Juan Angulo-Lozano, Daniel Ranti, Bernard H. Bochner, John P. Sfakianos, Amir Horowitz, Jeffrey V. Ravetch, David A. Knorr

**Author notes:** Correspondence: David Knorr, Jeffrey Ravetch.

## Abstract

CD40 is a central co-stimulatory receptor implicated in the development of productive anti-tumor immune responses across multiple cancers, including bladder cancer. Despite strong preclinical rationale, systemic administration of therapeutic agonistic antibodies targeting the CD40 pathway have demonstrated dose limiting toxicities with minimal clinical activity to date, emphasizing an important need for optimized CD40-targeted approaches, including rational combination therapy strategies. Here, we describe an important role for the endogenous IL-15 pathway in contributing to the therapeutic activity of CD40 agonism in orthotopic bladder tumors, with upregulation of trans-presented IL-15/IL-15Rα surface complexes, particularly by cross-presenting cDC1s, and associated enrichment of activated CD8 T cells within the bladder tumor microenvironment. In bladder cancer patient samples, we identify DCs as the primary source of IL-15, however, they lack high levels of IL-15Rα at baseline. Using humanized immunocompetent orthotopic bladder tumor models, we demonstrate the ability to therapeutically augment this interaction through combined treatment with anti-CD40 agonist antibodies and exogenous IL-15, including the fully-human Fc-optimized antibody 2141-V11 currently in clinical development for the treatment of bladder cancer. Combination therapy enhances the crosstalk between Batf3-dependent cDC1s and CD8 T cells, driving robust primary anti-tumor activity and further stimulating long-term systemic anti-tumor memory responses associated with circulating memory-phenotype T and NK cell populations. Collectively, these data reveal an important role for IL-15 in mediating anti-tumor CD40 agonist responses in bladder cancer and provide key proof-of-concept for combined use of Fc-optimized anti-CD40 agonist antibodies and agents targeting the IL-15 pathway. These data support expansion of ongoing clinical studies evaluating anti-CD40 agonist antibodies and IL-15-based approaches to evaluate combinations of these promising therapeutics for the treatment of patients with bladder cancer.

## Introduction

CD40, a member of the tumor necrosis factor superfamily of receptors (TNFRSF), plays a central co-stimulatory role in both innate and adaptive immunity, serving as a proximal regulator of the adaptive immune cascade driving the development of antigen-specific T cell responses (1). Evidence across multiple preclinical models for the ability of CD40 activation to induce robust anti-tumor responses (2), as well as the association of CD40 expression with improved clinical outcomes and markers of type-I immunity in pan-cancer human tumor analyses (3), have provided strong rationale for the clinical development of CD40 agonist approaches for cancer therapy, including agonistic anti-CD40 antibodies. Despite this promise, however, the clinical activity reported to date for these agents, particularly as systemic monotherapy, has been disappointing (2, 4), with transaminitis and thrombocytopenia reported at doses of 0.2 mg/kg, thus limiting the ability to achieve therapeutic doses for evaluation. These prior studies highlighted an important need for continued optimization of current CD40-targeted therapies.

We and others have previously implicated CD40 as an important pathway in the context of bladder cancer (5-9), a disease area of significant unmet clinical need with an estimated 550,000 new cases and nearly 200,000 disease-associated deaths annually worldwide (10). Using humanized orthotopic bladder cancer murine models, we have also recently identified the ability to substantially enhance therapeutic anti-tumor immune responses through the use of novel agonistic anti-CD40 antibodies engineered for optimal engagement of the FcγRIIB receptor essential for *in vivo* agonistic activity (5), an approach now being tested in an early-phase clinical study (NCT05126472) for the treatment of non-muscle invasive bladder cancer (NMIBC) unresponsive to front-line therapy. Here, we report on the mechanisms of the CD40 agonist response in preclinical bladder cancer models, with the identification of complementary pathways that provide a basis for rational combination therapy strategies for future clinical evaluation.

## Results

### Dendritic cells in the bladder microenvironment of mice responding to CD40 agonism have higher expression of IL-15Rα

As noted above, we previously identified CD40 as an important pathway in the immunoregulation of bladder cancer and further elucidated the ability of *in situ* CD40 agonism (via local intravesical treatment with agonistic anti-CD40 antibodies) to drive productive anti-tumor immunity in this disease context (5). While we previously found that anti-tumor immunity was critically dependent on conventional type 1 dendritic cells (cDC1s) and CD8 T cells, other signals participating in this response remain unknown. We hypothesized that IL-15 may be an important contributor to CD40 agonist therapeutic activity given the role of the IL-15 pathway in driving immune-stimulatory interactions between activated dendritic cells (DCs) and effector CD8 T cells and NK cells in multiple physiologic and pathologic contexts (11). To investigate this question in a bladder cancer-relevant setting, we utilized an immunocompetent orthotopic murine tumor model in which the syngeneic MB49 bladder cancer cell line is implanted into the bladders of C57BL/6J mice using a urethral catheter-based technique (**Fig. 1A**)(12, 13). This approach results in efficient engraftment of progressive bladder tumors recapitulating the enriched expression of CD40 within the bladder tumor microenvironment found in human disease (5, 6). We first evaluated for the presence of IL-15Rα on myeloid subsets in the TME of orthotopic MB49 bladder tumors in mice responsive versus unresponsive to intravesical CD40 agonism, as measured by luminescent signal at day 24 post-implantation (**Fig. 1B**). We found significantly higher amounts of cell surface expressed IL-15Ra on cDC1 and cDC2 subsets in responding tumors, without significant differences in expression on macrophages or neutrophils (**Fig. 1C**). These data demonstrate elevated IL-15Ra is associated with response to intravesical CD40 immunotherapy, suggesting that a direct role for IL-15/IL-15Ra may be driving the therapeutic effects of CD40 agonist antibodies in the bladder TME.

**Figure 1.**
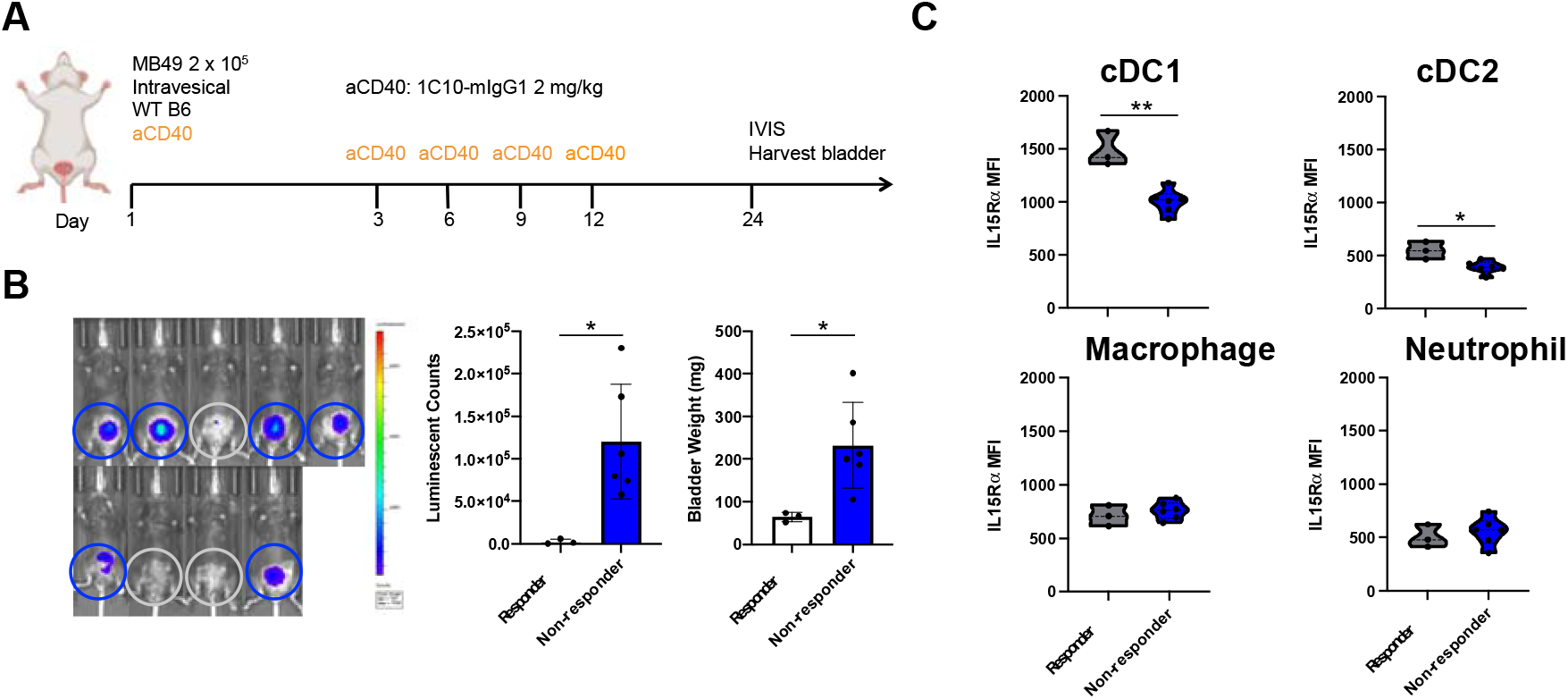
Dendritic cells in the bladder microenvironment of mice responding to CD40 agonism have higher expression of IL-15Rα. (*A*) Schematic of the treatment of mice bearing orthotopic MB49 bladder tumors with anti-CD40 antibody. (*B*) Representative intravital luciferase imaging (left) and quantification of luminescence (center) and bladder weights (right) across mice at day 24 post-tumor implantation (n = 3-6 mice per group; bars represent SD). Responders (gray circles) and non-responders (blue circles) to anti-CD40 antibody therapy are indicated. (*C*) IL-15Rα mean fluorescence intensity across mice on type-1 conventional DCs (cDC1; defined as F4/80^−^Ly-6G^−^CD11c^+^MHCII^+^XCR1^+^), type-2 conventional DCs (cDC2; defined as F4/80^−^Ly-6G^−^CD11c^+^MHCII^+^SIRPα^+^), macrophages (defined as CD11b^+^F4/80^+^Ly-6G^−^), and neutrophils (defined as CD11b^+^Ly-6G^+^) in the bladder microenvironment as assessed by flow cytometry at day 24 post-tumor implantation (n = 3-6 mice per group). *p < 0.05, **p < 0.01.

### Endogenous IL-15 participates in the therapeutic activity of CD40 agonism

To further evaluate the role of IL-15/IL-15Rα in the therapeutic response to CD40 agonism, mice bearing orthotopic bladder tumors were treated intravesically with a murine agonistic anti-CD40 antibody (IgG1 clone 1C10, capable of engaging murine FcγRIIB) either in the presence or absence of an IL-15 blocking antibody (**Fig. 2A**). Consistent with our prior study (14), intravesical anti-CD40 agonist antibody treatment results in decreased tumor burden, as assessed by both tumor cell bioluminescence and bladder weights (**Fig. 2B**). Notably, concurrent IL-15 blockade was associated with a reduced therapeutic effect in response to anti-CD40 agonist antibody treatment, suggesting that endogenous IL-15 participates at least in part in the therapeutic activity of CD40 agonism in this tumor model.

**Figure 2.**
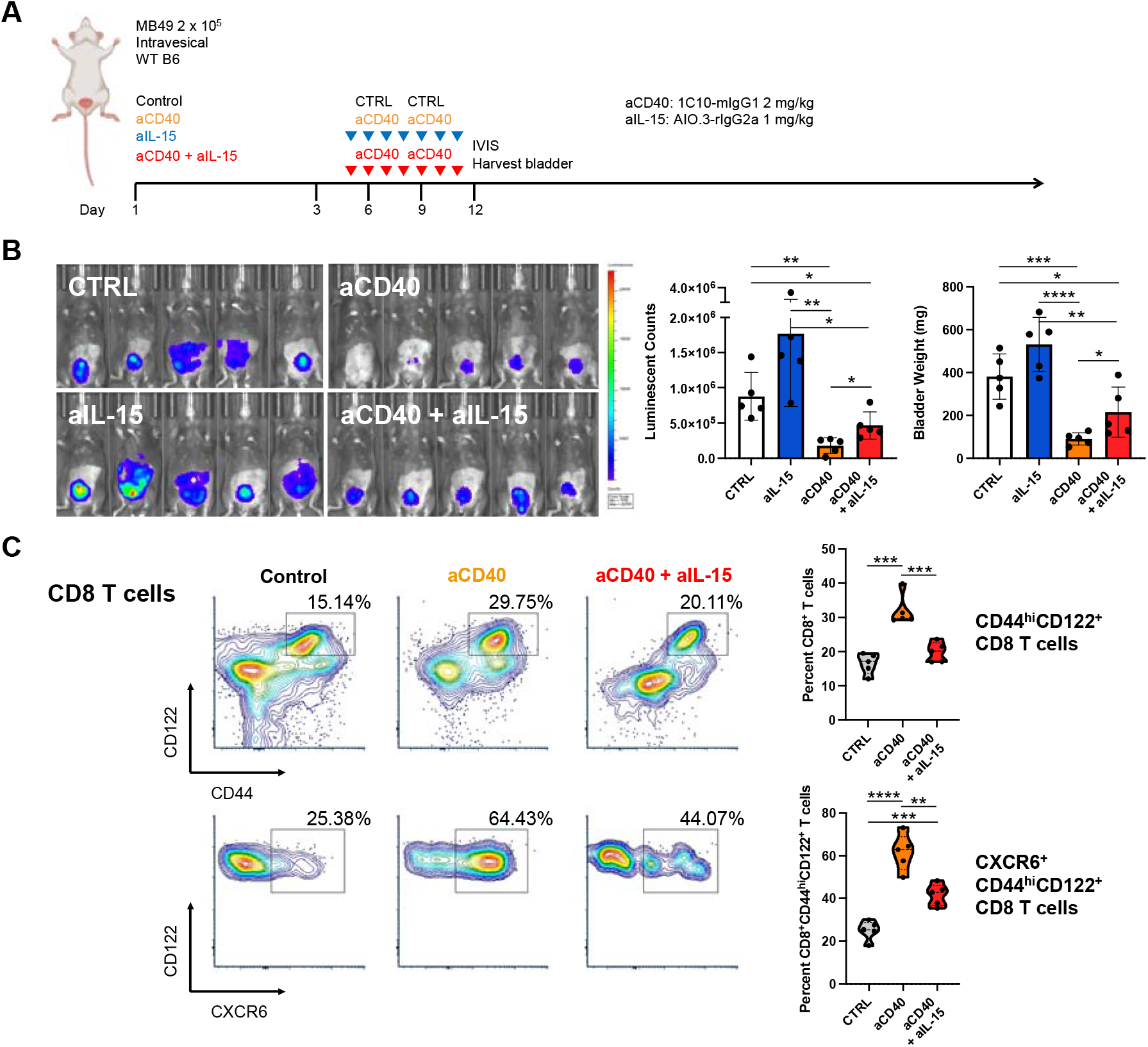
Endogenous IL-15 participates in the therapeutic activity of CD40 agonism. (*A*) Schematic of the treatment of mice bearing orthotopic MB49 bladder tumors with anti-CD40 antibody and/or anti-IL-15 blocking antibody or isotype-matched control antibody. (*B*) Representative intravital luciferase imaging (left) and quantification of luminescence (center) and bladder weights (right) across mice at day 12 post-tumor implantation (n = 5 mice per group; bars represent SD). (*C*) Representative flow cytometry plots and quantification across mice (n = 5 mice per group) of CD44^hi^CD122^+^ CD8 T cells (top) and CXCR6^+^CD44^hi^CD122^+^ CD8 T cells (bottom) in the bladders of mice treated as outlined in A. *p < 0.05, **p < 0.01, ***p < 0.001, ****p < 0.0001.

We have previously demonstrated that productive responses to CD40 agonist therapy in the orthotopic bladder tumor setting is CD8 T cell-dependent (5, 15), thus we further examined the tumor-infiltrating CD8 T cell compartment. Prior work has demonstrated that antigen-specific CD8 T cells expressing the IL-2R*β* chain (CD122) are potently activated when seeing IL-15 *in trans* from DCs in the TME, and their positioning to this niche is driven by the chemokine receptor CXCR6 (16). We found the therapeutic activity induced by anti-CD40 agonist antibody treatment was associated with increased proportions of activated CD44^hi^CD122^+^ CD8 T cells in the bladder microenvironment, which was reduced in the setting of concurrent IL-15 blockade (**Fig. 2C, top**). Interestingly, anti-CD40 agonist antibody treatment was also associated with increased proportions of activated CD8 T cells expressing CXCR6 (**Fig. 2C, bottom**), further supporting its important role in optimizing intratumoral IL-15-driven interactions between DCs and effector T cells necessary for sustained tumor control.

### Dendritic cells in the TME of non-muscle invasive bladder cancer patients express high levels of CD40 and IL-15 but are limited in their expression of IL-15Rα

To determine whether similar pathways in dendritic cells may be operative in patients with bladder cancer, we next used single cell RNA sequencing (scRNAseq) to evaluate patients with NMIBC who were untreated (n=3) or from NMIBC patients that were identified to be BCG unresponsive (n=5). Cells were filtered according to minimum gene count (n=200); genes were filtered according to minimum expression totals (n=3); cells with high mitochondrial gene counts (>20%) were filtered out; samples were normalized and log-transformed; and all samples were concatenated together. A total of 28,324 cells were isolated in the final population of this analysis. 6,921 tumor cells; 6,391 CD4+ T cells; 3,714 B cells; 2,621 monocytes/macrophages; 2,641 CD8+ T cells; 1756 stromal cells; 1281 NK cells; 1144 fibroblasts; 23 plasma cells; 563 endothelial cells; 145 mature DCs; and 581 unknown cells were labelled. We first used canonical lineage markers for cell populations were utilized to assign cell identities to each cluster (**Fig. 3A**). tSNE plots were created for four representative markers: CD40, CD40L, IL15, and IL15RA (**Fig. 3B**). Of the scaled expression values across patients, these data confirmed that dendritic cells were among the highest expressors of both CD40 and IL-15 (**Fig. 3C**). However, at baseline DCs did not demonstrate increased levels of IL-15RA, which was higher in endothelial cells and plasma cells, as previously demonstrated(17). Notably, T cells were the highest expressors of CD40 ligand (CD40L, CD154), representing their known role for their ability to provide T cell help to DCs and B cells. These data confirm that while DCs of the NMIBC TME express high levels of CD40 and IL-15, the expression IL-15RA, which is necessary for optimal trans-presentation to effector CD8 T cells, is limited.

**Figure 3.**
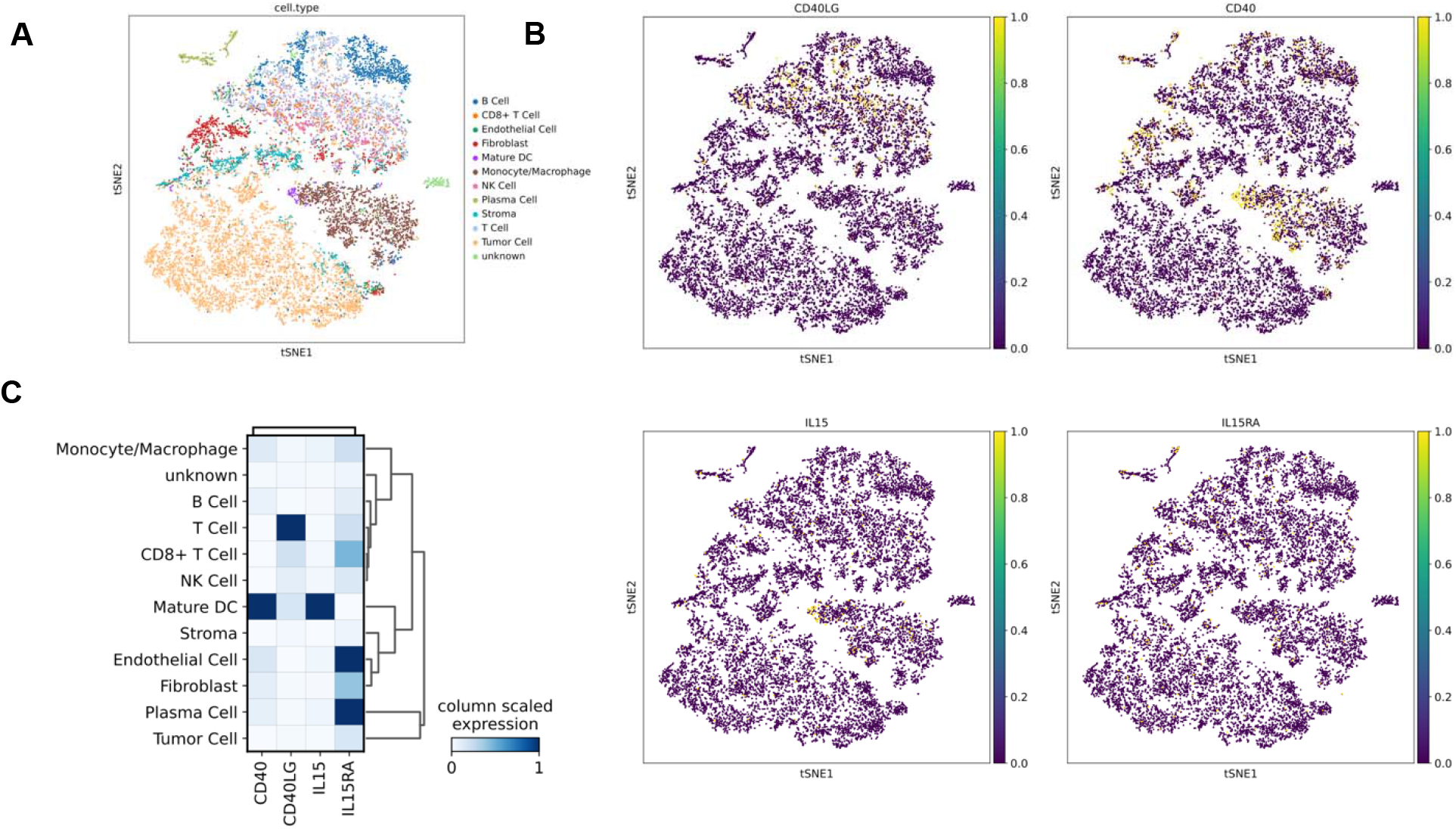
DCs in human non-muscle invasive bladder cancer express high levels of IL-15. Eight primary tumor samples from either untreated NMIBC patients (n=3) or from NMIBC patients that had progressed from NMIBC to MIBC following BCG therapy (n=5) were included in this single-cell RNA sequencing analysis. A) Phenograph clustering was performed and 29 clusters were identified. Canonical lineage markers for cell populations were utilized to assign cell identities to each cluster. B) tSNE plots were created for four representative markers: CD40, CD40L, IL15, and IL15RA. C) A matrix plot with column scaled expression of the same four markers of interest is shown to highlight the distribution and intensity of expression for each gene across each subset.

### IL-15/IL-15Rα is upregulated on dendritic cells in the bladder tumor microenvironment in response to CD40 agonism

IL-15 is known to be co-expressed *in trans* with IL-15Rα by myeloid cells, including by DCs, where IL-15Rα binds to IL-15 intracellularly and functions as a chaperone protein on the cell surface (18-23). Surface IL-15/IL-15Rα complexes are trans-presented to opposing lymphocytes during cell-cell interactions are primary mechanism by which physiologic IL-15 signals are delivered *in vivo* (24). We therefore investigated the expression of IL-15Rα in our orthotopic bladder tumor models. We observed that intravesical anti-CD40 agonist antibody treatment leads to upregulation of IL-15Rα on DCs and other myeloid cells within the bladder tumor microenvironment (**Fig. 4A and B**). Notably, the highest expression of IL-15Rα was found in the DC compartment, the population that has been likewise identified to be the highest expressors of CD40 in the bladder tumor context (5). Moreover, the cDC1 subset, previously implicated as the key antigen presenting cell (APC) in the bladder tumor microenvironment essential for cross-presenting antigen to CD8 T cells and driving productive responses to CD40 agonist therapy (5), was identified to have the highest expression of IL-15Rα. Of note, IL-15Rα upregulation was not similarly observed upon treatment with BCG (**Fig. 4A and B**), a live-attenuated bacterium routinely used clinically as a therapeutic immune stimulus in the treatment of localized bladder cancer, supporting the potential unique cooperation of the CD40 and IL-15 pathways in the bladder tumor microenvironment.

**Figure 4.**
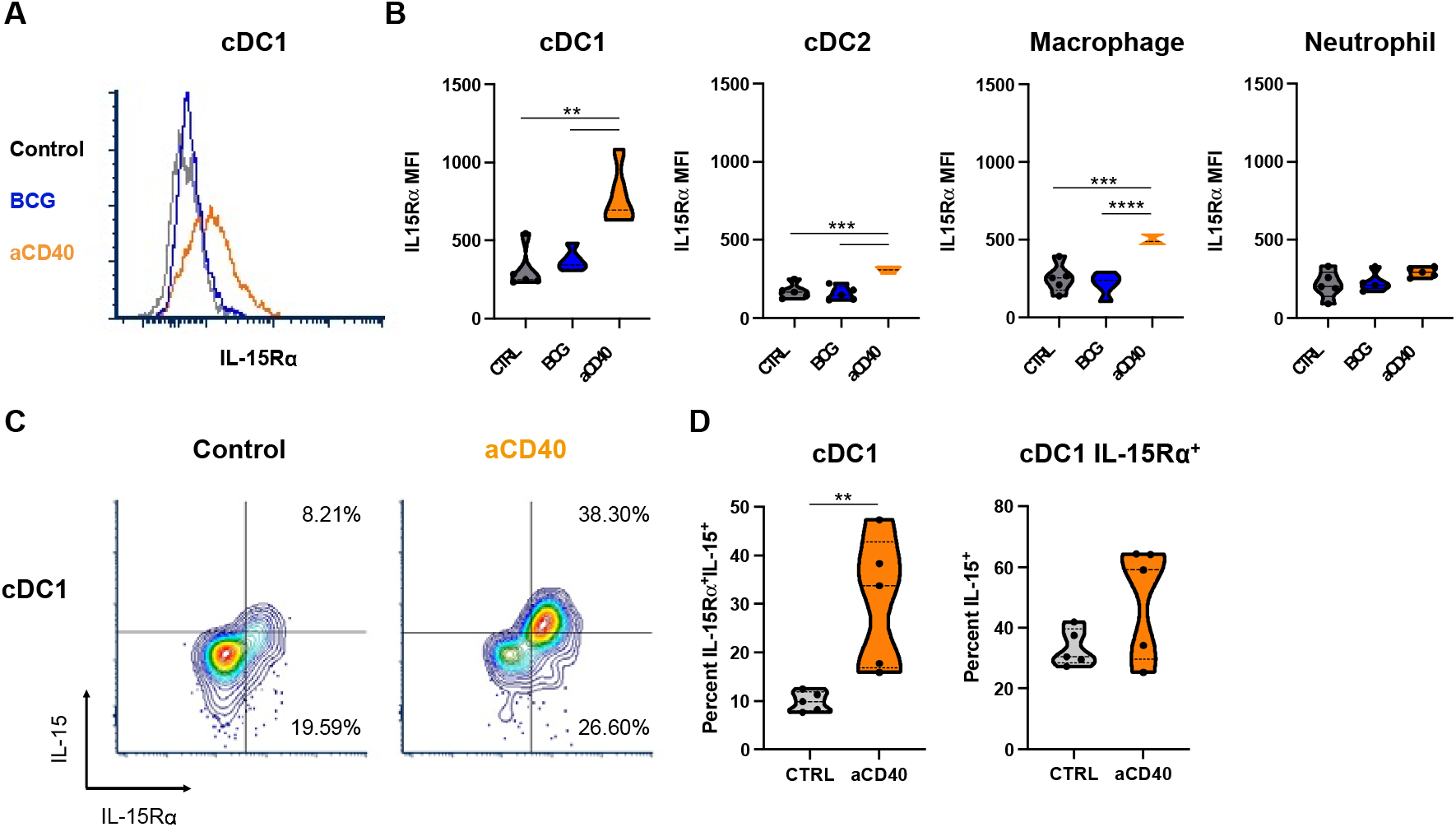
CD40 agonism induces IL-15/IL-15Rα upregulation on dendritic cells in the bladder tumor microenvironment. Mice bearing orthotopic MB49 bladder tumors were treated intravesically with anti-CD40 antibody, BCG, or isotype-matched control antibody on days 6 and 9 post-tumor implantation. (*A*) Representative histogram and (*B*) quantification across mice of IL-15Rα mean fluorescence intensity on type-1 conventional DCs (cDC1; defined as F4/80^−^Ly-6G^−^CD11c^+^MHCII^+^XCR1^+^), type-2 conventional DCs (cDC2; defined as F4/80^−^Ly-6G^−^CD11c^+^MHCII^+^SIRPα^+^), macrophages (defined as CD11b^+^F4/80^+^Ly-6G^−^), and neutrophils (defined as CD11b^+^Ly-6G^+^) in the bladder microenvironment as assessed by flow cytometry at day 12 post-tumor implantation (n = 5 mice per group). (*C*) Representative flow cytometry plots and (*D*) quantification across mice of IL-15 and IL-15Rα surface expression on cDC1s in the bladder microenvironment as assessed by flow cytometry at day 12 post-tumor implantation (n = 5 mice per group). Left graph is gated on cDC1s and depicts proportions of cDC1s that are double-positive for surface IL-15 and IL-15Rα. Right graph is gated on IL-15Rα-expressing cDC1s and depicts proportions of cDC1s expressing IL-15Rα that is occupied by IL-15. **p < 0.01, ***p < 0.001, ****p < 0.0001.

To corroborate our findings in the MB49 model, we examined an additional immunocompetent orthotopic bladder cancer model derived from the syngeneic UPPL1541 bladder tumor cell line, a cell line generated from a genetically engineered murine model of bladder cancer (*Upk3a-Cre*^*ERT2*^; *Trp53*^*L/L*^; *Pten*^*L/L*^; *Rosa26*^*LSL-Luc*^) that recapitulates the luminal molecular subtype of human high-grade urothelial carcinoma (25). IL-15Rα upregulation in response to CD40 agonism was similarly observed in the setting of mice bearing orthotopic UPPL1541 tumors (**SI Appendix, Fig. S1**). These observations across two immunocompetent bladder tumor models suggest that CD40 agonist-driven upregulation of the IL-15 pathway may be a broader phenomenon in the bladder tumor microenvironment.

IL-15Rα protein expression has been previously postulated to likely exceed that of IL-15, given readily detectable IL-15Rα on a range of cell types and much more limited detection of surface-associated IL-15 (26). Concurrent surface staining of IL-15Rα and IL-15 revealed upregulation of surface IL-15/IL-15Rα complexes on cDC1s in the bladder microenvironment in response to CD40 agonism (**Fig. 4C and D**). However, a notable proportion of the IL-15Rα expressed on the cell surface was not found to be occupied by IL-15, both at baseline and following CD40 agonist therapy (**Fig. 4D, bottom right quadrant**).

### Combination therapy with a fully-human Fc-optimized anti-CD40 agonist antibody and IL-15 enhances primary anti-tumor activity in humanized mouse models of bladder cancer

The above data and the prior literature (18-23) thus support the hypothesis that exogeneous IL-15 might provide an opportunity to further enhance CD40 agonist therapeutic activity. We tested this hypothesis using a CD40 and FcyR humanized C57BL6 mouse and the fully-human anti-CD40 agonist antibody 2141-V11, an antibody Fc-engineered for enhanced FcγRIIB binding necessary for optimal CD40 agonist activity (27) that is under active clinical evaluation for the intravesical treatment of NMIBC (NCT05126472). The humanized hCD40/hFcγR model recapitulates the expression patterns and function of human CD40 and human FcγR to allow full *in vivo* assessment of the interaction of fully-human anti-CD40 antibodies within the unique human FcγR landscape, necessary for accurate evaluation of the *in vivo* activity of these antibodies (15, 27).

Using this humanized immunocompetent hCD40/hFcγR *in vivo* platform, we examined the therapeutic activity of the human anti-CD40 agonist antibody 2141-V11 either alone or in combination with IL-15 against orthotopic MB49 bladder tumors (**Fig. 5A**). In this setting, the combination of intravesical 2141-V11-driven CD40 agonism and systemic IL-15 was found to substantially decrease tumor burden assessed at earlier time points by tumor cell bioluminescence (**Fig. 5B**), as well as significantly improve rates of complete and durable response and long-term overall survival (**Fig. 5C**). We similarly observed improved anti-tumor activity against orthotopic UPPL1541 bladder tumors in response to combined therapy with the anti-CD40 agonist antibody 2141-V11 and IL-15, as visualized by serial bladder ultrasound and the time to tumor detection (**SI Appendix, Fig. S2**). These data indicate the capability of this combination in promoting strong primary anti-tumor activity across immunocompetent orthotopic bladder cancer models. Of note, while CD40 agonist therapy results in robust induction of IL-15/IL-15Rα (**Fig. 2 and SI Appendix, Fig. S2**), exogenous IL-15 therapy was not observed to reciprocally enhance CD40 expression by any of the examined CD40-expressing myeloid cell populations within the bladder tumor microenvironment (**SI Appendix, Fig. S3**).

**Figure 5.**
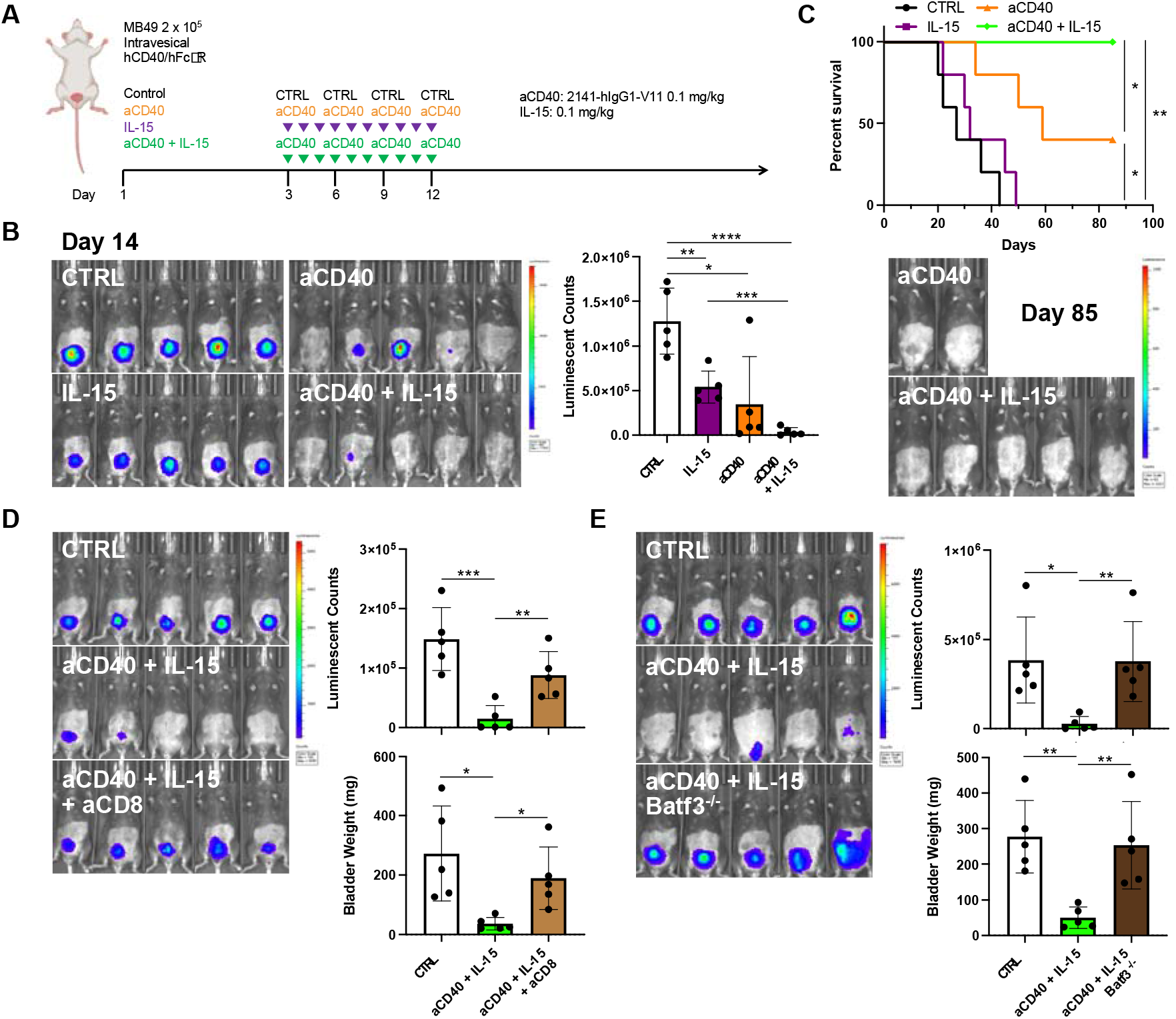
Combination therapy with Fc-optimized anti-CD40 agonist antibody 2141-V11 and IL-15 enhances primary anti-tumor activity. (*A*) Schematic of the treatment of humanized hCD40/hFcyR mice bearing orthotopic MB49 bladder tumors with anti-CD40 antibody 2141-V11 and/or IL-15 or control (isotype-matched control antibody and/or vehicle). (*B*) Representative intravital luciferase imaging (left) and luminescence quantification (right) across mice at day 14 post-tumor implantation (n = 5 mice per group; bars represent SD). (*C*) Survival (top) and representative intravital luciferase imaging (bottom) of surviving mice at day 85 post-tumor implantation treated as outlined in A. (*D*) Representative intravital luciferase imaging (left) and quantification of luminescence (right, top) and bladder weights (right, bottom) at day 14 post-tumor implantation across mice treated with anti-CD40 antibody 2141-V11 and IL-15 or control in the absence or presence of additional anti-CD8 T cell depleting antibody (n = 5 mice per group; bars represent SD). (*E*) Representative intravital luciferase imaging (left) and quantification of luminescence (right, top) and bladder weights (right, bottom) at day 14 post-tumor implantation across WT or Batf3^-/-^ mice treated with anti-CD40 antibody 1C10 and IL-15 or control (n = 5 mice per group; bars represent SD). *p < 0.05, **p < 0.01, ***p < 0.001.

### CD8 T cells and Batf3 are required for the anti-tumor activity of combined therapeutic targeting of CD40 and IL-15

We further evaluated the cellular mediators by which combined therapeutic activation of CD40 and IL-15 results in anti-tumor responses in the bladder tumor context. Depletion of CD8 T cells using an anti-CD8 depleting antibody largely abrogated the anti-tumor activity of the therapeutic combination, as assessed by tumor cell bioluminescence and bladder weights (**Fig. 5D**). This suggests that the therapeutic response to combined CD40 agonist and IL-15 targeting requires CD8 T cells in this setting.

Given the central role of cDC1s in mediating responses to CD40 agonist monotherapy (5) and the high expression of IL-15Rα induced in this population by CD40 agonism within the bladder tumor microenvironment (**Fig. 4 and SI Appendix, Fig. S1**), we hypothesized that cDC1s were likewise important in driving the anti-tumor activity of CD40/IL-15-targeted combination therapy. We and others have previously shown that Batf3^-/-^ mice are deficient in cDC1s, while other populations, including cDC2s and macrophages, remain intact (5, 28, 29). We observed that the reduced bladder tumor burden driven by combination therapy with anti-CD40 agonist antibody (1C10) and IL-15 was lost in the Batf3^-/-^ background (**Fig. 5E**), indicating a requirement for Batf3 in the therapeutic activity of this combination. Taken together, these data support the conclusion that the cDC1-CD8 T cell axis, impaired in the Batf3^-/-^ setting, plays a critical role in driving the response to combined targeting of the CD40 and IL-15 pathways in this context.

### Combined therapeutic targeting of CD40 and IL-15 promotes systemic anti-tumor memory responses

Prior studies have described an important role for IL-15 in supporting the induction and maintenance of immune effector cell memory responses (21, 26, 30, 31). Using the humanized hCD40/hFcγR *in vivo* model, we further interrogated the ability of combined treatment with the anti-CD40 agonist antibody 2141-V11 and IL-15 to induce long-term protective systemic immunity. Mice surviving long-term (>90 days) from the initial therapy were rechallenged at a ten-fold higher dose of tumor cells in a different tissue compartment (subcutaneous rather than within the bladder) in the absence of any additional therapy (**Fig. 6A**). Mice previously treated with 2141-V11-driven CD40 agonism and IL-15 uniformly rejected tumor rechallenge (**Fig. 6B**), indicating that this treatment is capable of inducing an *in situ* vaccination effect driving durable protective systemic anti-tumor immunity.

**Figure 6.**
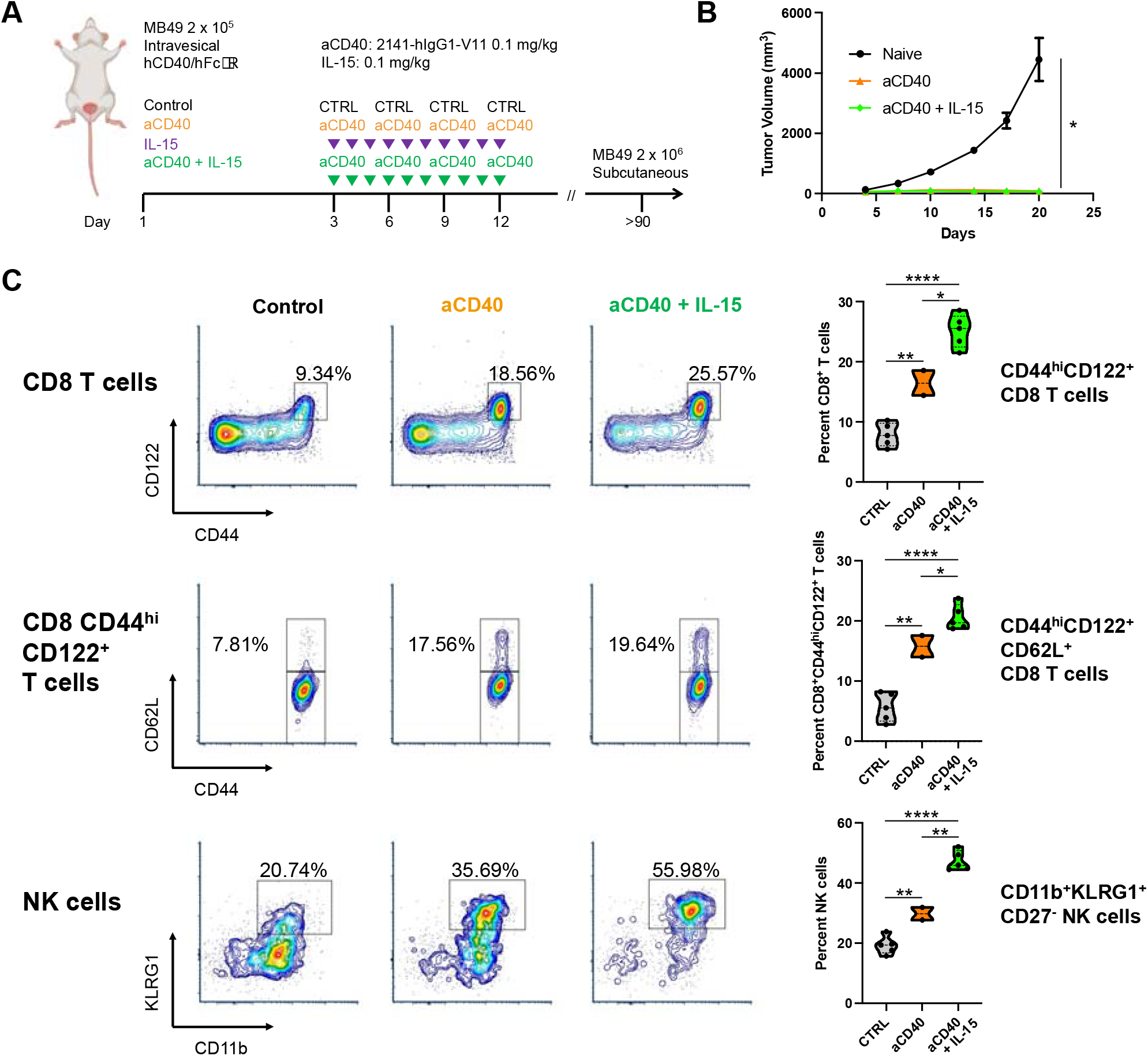
Combination therapy with Fc-optimized anti-CD40 agonist antibody 2141-V11 and IL-15 enhances anti-tumor memory responses. (*A*) Schematic of subcutaneous rechallenge (ten-fold tumor cell dose) of humanized hCD40/hFcyR mice surviving long-term (>90 days after initial orthotopic MB49 bladder tumor implantation) following initial therapy with anti-CD40 antibody 2141-V11 with or without IL-15. (*B*) Tumor growth (in the absence of any additional therapy) of long-term survivors compared with naïve mice receiving an equivalent MB49 tumor cell implant (n = 2-5 mice per group; bars represent SD). (*C*) Representative flow cytometry plots and quantification across mice (n = 2-5 mice per group) of CD44^hi^CD122^+^ CD8 T cells (top), CD44^hi^CD122^+^CD62L^+^ CD8 T cells (middle), and CD11b^+^KLRG1^+^CD27^−^ NK cells (bottom) in the peripheral blood of mice surviving long-term (day 110) after primary tumor implantation and tumor rechallenge. *p<0.05, **p<0.01, ****p<0.0001.

Given this evidence of a robust systemic memory response, we further examined lymphocytes in the peripheral blood of these mice surviving long-term after both primary tumor treatment and subsequent tumor rechallenge to assess potential phenotypic changes associated with this response. Mice initially treated with the anti-CD40 agonist antibody 2141-V11 and IL-15 demonstrated increased proportions of circulating CD44^hi^CD122^+^CD62L^+^ CD8 T cells (**Fig. 6*C, top and middle***), consistent with enrichment of CD8 T cells bearing a central memory phenotype. Combination therapy targeting CD40 and IL-15 was also associated with increased proportions of circulating, CD27^−^CD11b^+^KLRG1^+^ NK cells (**Fig. 6*C, bottom***), a phenotype associated with a population of mature NK cells that has been implicated in contributing to durable immune responses in several contexts (32). Together, these data suggest that combined therapeutic targeting of the CD40 and IL-15 pathways results in enrichment of memory-phenotype CD8 T cell and NK cell populations, which correlate with long-term enhanced systemic anti-tumor memory responses.

## Discussion

Our group has previously described the importance of FcyRIIB engagement for optimal in vivo activity of anti-CD40 agonistic antibodies (5, 27). While the potential of this Fc-optimized approach to improve activity in patients is encouraging, optimal clinical benefit is most likely to be achieved through additional rationally selected combination(33). We thus set out to define such pathways in the context of therapeutic CD40 agonism and establish proof-of-concept for potential concurrent therapeutic targeting with direct implications for clinical translation.

Here, we identify an important role for the endogenous IL-15 pathway in contributing to the therapeutic activity of CD40 agonism in the bladder TME, associated with the interaction between cross-presenting Batf3-dependent cDC1s and CD8 T cells, as well as trans-presented IL-15/IL-15Rα surface complexes. DCs, in particular cDC1s, have been increasingly implicated as a critical component of cancer immune surveillance and effective immunotherapy responses (34-37), with cytokines such as IFNγ and IL-12 previously described to play important roles in the productive interaction between DCs and T cells driven by CD40 activation in the tumor microenvironment (38, 39). The current study identifies IL-15 as a key additional signal in the bladder cancer context. The substantial (but not complete) abrogation of CD40 agonist-driven anti-tumor activity associated with IL-15 blockade suggests a mechanism involving multiple signals including, but not limited to, IL-15, consistent with the likely additional involvement of IL-12 and IFNγ discussed above. Indeed, cooperation between IL-12, IL-15, and IFNγ in orchestrating protective immunity has been suggested in cancer and other contexts (22, 40, 41). Di Pilato et al has also recently observed a critical role for a population of intratumoral IL-12-expressing DCs in mediating the survival and local expansion of effector-like CD8 T cells via an IL-15 trans-presentation mechanism, found to be necessary for sustained tumor control and orchestrated in perivascular niches within the tumor microenvironment through the CXCR6-CXCL16 chemotactic axis (16). In the present study, we observed an IL-15-dependent enrichment of activated CXCR6-expressing CD8 T cells in the bladder tumor microenvironment following CD40 agonist therapy. Interestingly, RNA sequencing analysis of patient tumor specimens has also recently revealed a correlation between the expression of CD40 and CXCR6 across multiple human tumor types (4). Our own sequencing data demonstrated that indeed DCs in the NMIBC TME are the highest expressors of CD40 and IL-15, however, they do not express high levels of IL-15RA. It is intriguing to consider whether the anti-tumor activity of therapeutic CD40 activation might be mediated in significant part through augmentation of this specific DC-T cell axis involving IL-15, IL-12, and CXCR6, which may occur spontaneously in certain immunogenic tumors (16), but is likely to be at least suboptimal in many other cancer settings like NMIBC. Whether the activity of DCs in the present context occurs primarily within the intratumoral microenvironment or at the level of the tumor-draining lymph nodes is an additional question of interest, one that may have important therapeutic implications, for instance, in selecting optimal routes and sequencing of therapeutic delivery.

Notably, in this study we further identify the opportunity to augment this productive anti-tumor DC-T cell interaction through combined therapeutic targeting of the CD40 and IL-15 pathways. Using two humanized immunocompetent orthotopic bladder tumor models and a clinically-relevant investigational approach, including use of the fully-human, Fc-optimized anti-CD40 agonist antibody 2141-V11 that is under active clinical evaluation for the treatment of bladder cancer and other indications (NCT05126472, NCT04059588, NCT04547777), we noted substantial improvement in the therapeutic activity of 2141-V11-induced CD40 agonism through concurrent stimulation with exogenous IL-15, including robust generation of both primary and systemic memory responses. This combination therapy approach was found to critically depend on an intact cDC1-CD8 T cell axis, and is likely to be leveraging the observed upregulation (but incomplete occupancy) of surface IL-15Rα, which has been shown to be capable of incorporating exogenous IL-15 into the trans-presentation pathway (42, 43). Other groups have also observed the ability of CD40 agonists to overcome SOCS3-mediated impairment of CD4 T cell help induced by immunotherapy with common cytokine receptor gamma chain (γc) family members like IL-15 (44), providing additional mechanistic support for the combined therapeutic activation of the CD40 and IL-15 pathways. While the therapeutic activity was found to be primarily CD8 T cell-dependent in this setting, notable pharmacodynamic changes were also observed in the NK cell compartment, consistent with the enhancement of effector lymphocyte activity likely to be broadly relevant across tumor contexts (45-47).

In recent years, IL-15 has gained considerable interest as a cancer immunotherapy target given its potent physiologic role in promoting the activation, proliferation, survival, cytotoxicity, and other functions of CD8 T cells and NK cells, including the development and maintenance of memory CD8 T cell and NK cell populations (11, 48). Indeed, deficits in IL-15 expression have been associated with worse clinical outcomes and reduced intratumoral immune infiltrates in analysis of patient tumor cohorts (49). In addition, in contrast to the related common γc cytokine IL-2, IL-15 has not been similarly implicated in mediating activation-induced cell death of effector T cells or promoting the activity of regulatory T cells (50-53). Multiple IL-15-based therapeutic formats are currently under active clinical evaluation for cancer therapy, including monomeric formulations modified for improved pharmacokinetics and biologic activity, as well as several heterodimeric approaches incorporating partial or full-length IL-15Rα, mimicking the biologically-active IL-15/IL-15Rα complexes found physiologically *in vivo* (54, 55).

Of particular interest in the bladder cancer setting is N-803 (formerly ALT-803), an IL-15 superagonist complex consisting of an IL-15 mutein (N72D) associated with a dimeric IL-15Rα sushi domain/IgG1 Fc fusion protein (56). N-803 in combination with BCG is currently being investigated in the phase 2/3 QUILT 3.032 study (NCT03022825) for the treatment of patients with NMIBC unresponsive to front-line BCG therapy. Updated clinical results for 160 patients treated with N-803 + BCG [83 patients with carcinoma in situ (CIS); 77 patients with papillary-only disease] were recently presented, reporting a 71% complete response rate with a 26.6 month median duration of response in patients with CIS, as well as a 51% disease-free survival rate at 18 months in patients with papillary-only disease (57). These results compare favorably to the outcomes for patients with BCG-unresponsive disease receiving historical salvage therapies (58, 59), as well as outcomes from the registrational phase 2 KEYNOTE-057 study of the PD-1 inhibitor pembrolizumab, which was recently-approved for treatment of the BCG-unresponsive CIS patient population (60). While these clinical data provide strong evidence for the therapeutic utility of IL-15 pathway activation in bladder cancer, continued rates of non-response and disease recurrence despite this therapy highlight an ongoing need for improved approaches. In this setting, we propose further investigation of novel IL-15-based combinations incorporating optimized CD40 agonists. Building on our current work establishing preclinical proof-of-concept for concurrent therapeutic targeting of CD40 and IL-15 in the bladder cancer context, as well as our ongoing phase 1 evaluation of the Fc-enhanced anti-CD40 agonist antibody 2141-V11 in patients with BCG-unresponsive NMIBC (NCT05126472), our group is now actively assessing specific IL-15-based therapeutic formats that may optimally synergize with 2141-V11-driven CD40 agonism to directly inform the near-term development of combination therapy clinical trials for this patient population.

Collectively, these data reveal an important role for IL-15 in mediating anti-tumor CD40 agonist responses in bladder cancer and provide key proof-of-concept for a rationally designed immune-based therapeutic approach combining use of Fc-optimized anti-CD40 agonist antibodies and agents targeting the IL-15 pathway, capable of driving enhanced cDC1 and CD8 T cell interaction and robust primary and memory anti-tumor responses. These data provide a rationale for expansion of ongoing clinical studies evaluating the Fc-optimized CD40 agonist antibody 2141-V11 to include novel IL-15-based combinations for the treatment of patients with bladder cancer.

## Materials and Methods

### Mice

C57BL/6J (WT; RRID:IMSR_JAX:000664) and B6.129S(C)-*Batf3*^*tm1Kmm*^/J (*Batf3*^*-/-*^; RRID:IMSR_JAX:013755) mice were purchase from The Jackson Laboratory. Mice expressing human *CD40* and human *FCGR1A, FCGR2A*^R131^, *FCGR2B*^I232^, *FCGR3A*^F158^, and *FCGR3B* under control of their endogenous human regulatory elements on an isogenic background deleted for the homologous mouse genes were generated and extensively characterized as previously described (27, 61). All mice were 8-12 weeks of age at the time of experimental use, and were bred and/or maintained in the Rockefeller University Comparative Bioscience Center under specific pathogen-free conditions. All experiments were performed in compliance with institutional guidelines and applicable federal regulations under protocols (17026-H, 20029-H) approved by the Rockfeller University Institutional Animal Care and Use Committee.

### Cell lines

Syngeneic murine bladder tumor cell lines MB49-luciferase (M. Glickman, Memorial Sloan Kettering) and UPPL1541 (W. Kim, University of North Carolina) were cultured in vacuum-gas plasma-treated tissue culture flasks (Falcon) at 37°C and 5% CO_2_ and maintained in Dulbecco’s Modified Eagle Medium (DMEM; Gibco) supplemented with 10% fetal bovine serum (Sigma), 100 U/mL penicillin (Gibco), and 100 μg/mL streptomycin (Gibco).

### Antibodies

Anti-human CD40 antibody 2141-V11 (derived from CP-870,893, clone 21.4.1 referenced in patent US7338660, ATCC accession number PTA-3605) containing a human IgG1 Fc domain carrying the G237D/P238D/H268D/P271G/A330R amino acid modifications was generated as previously described (27). Anti-mouse CD40 antibody 1C10 containing a mouse IgG1 Fc domain was generated as previously described (62). Plasmid sequences were validated by direct sequencing (Genewiz). Antibodies were produced by transient co-transfection of Expi293F cells (maintained in serum-free Expi293 Expression Medium) with heavy- and light-chain constructs using the ExpiFectamine 293 Transfection Kit (Thermo Fisher Scientific), and subsequently purified using Protein G Sepharose 4 Fast Flow (GE Healthcare), eluted using IgG elution buffer (Thermo Fisher Scientific), dialyzed in PBS, and sterile-filtered, as previously described (63). Anti-mouse IL-15 (clone AIO.3) and anti-mouse CD8α (clone 2.43) depleting antibodies, as well as all isotype control antibodies, were purchased from Bio X Cell.

### Intravesical tumor implantation and treatments

Tumor cells were implanted orthotopically into the bladders of mice using a catheter-based intravesical protocol, as previously described (5). In brief, tumor cells were harvested using 0.05% trypsin-EDTA (Gibco) for 5 min at 37°C, washed twice with DMEM, assessed for cell count and viability via trypan blue staining (Millipore) using a Countess II automated cell counter (Thermo Fisher Scientific), and resuspended in DMEM (4 × 10^6^ cells/mL for MB49-luciferase; 2 × 10^8^ cells/mL for UPPL1541). Mice were anesthetized with inhalational isoflurane, bladders were voided with digital pressure, and a polyurethane 24G x 3/4” catheter (Terumo) was inserted transurethrally into the bladder for intravesical instillation. Prior to tumor cell instillation, protamine sulfate (10 mg/mL in H_2_O; Sigma) was instilled into the bladder (50 μL) and maintained for 30 min, and subsequently voided by digital pressure. Tumor cell suspension was then instilled into the bladder (50 μL) and maintained for 60 min. Heating pads were used to maintain core body temperature throughout implantation and recovery. For intravesical treatment, mice were anesthetized with inhalational isoflurane, bladders were voided with digital pressure, and a polyurethane 24G x 3/4” catheter was inserted transurethrally into the bladder. Treatments were then instilled into the bladder (anti-human CD40 antibody 2141-V11 at 2.5 μg in 50 μL per dose; anti-mouse CD40 antibody 1C10 at 50 μg in 50 μL per dose; BCG TICE strain at 4 × 10^6^ colony-forming units in 50 μL per dose) and maintained for 60 min (see figures/figure legends for individual experimental treatment schemas). When indicated, the following agents were administered via intraperitoneal injection: recombinant mouse IL-15 (2.5 μg/dose; PeproTech), anti-mouse IL-15 antibody (25 μg/dose; clone AIO.3), and anti-mouse CD8α antibody (250 μg/dose; clone 2.43).

### Subcutaneous tumor rechallenge

Mice surviving greater than 90 days after primary treatment of orthotopically-implanted MB49-luciferase bladder tumors were confirmed to be free of detectable tumor by bioluminescence imaging. Cultured MB49-luciferase tumor cells were harvested using 0.05% trypsin-EDTA for 5 min at 37°C, washed twice with DMEM, assessed for cell count and viability, and resuspended in DMEM (4 × 10^7^ cells/mL). Mice were injected subcutaneously in their lower flank with 50 μL of the tumor cell suspension. Tumors were subsequently measured 2-3 times per week using an electronic caliper and tumor volume was calculated using the formula (L_1_ ^2^ × L_2_)/2, with L_1_ and L_2_ corresponding to the shortest and longest dimensions, respectively.

### Bioluminescence imaging

Mice bearing orthotopic MB49-luciferase tumors were administered 3 mg of D-luciferin bioluminescent substrate (PerkinElmer) via intraperitoneal injection. Prior to imaging, mice were anesthetized with inhalational isoflurane and abdominal hair was removed via shaving. Images were subsequently acquired 20 min after luciferin injection using an IVIS Spectrum *in vivo* imaging system (PerkinElmer) with a 10 s exposure time. Total luminescence counts were quantified across a standardized region of interest centered on the bladder area using Living Image analysis software (PerkinElmer).

### Ultrasound imaging

Prior to imaging, mice were anesthetized with inhalational isoflurane, abdominal hair was removed via shaving and application of chemical hair removal cream, and ultrasound gel was applied to the abdomen. Bladders were subsequently imaged using a Vevo 2100 ultrasound imaging system (FUJIFILM VisualSonics). Bladders were monitored by ultrasound twice per week beginning on day 8 after tumor implantation to assess for the appearance of bladder tumors.

### Flow cytometry

Tumor and bladder tissue were harvested and processed into single-cell suspensions using the Mouse Tumor Dissociation Kit and gentleMACS Octo Dissociator with Heaters (Miltenyi Biotec) according to the manufacturer’s protocols. Peripheral blood was collected via retro-orbital sinus sampling from mice anesthetized with inhalational isoflurane and red blood cells were lysed with RBC Lysis Buffer (BioLegend) according to the manufacturer’s protocols. Cells were stained for viability using the LIVE/DEAD Fixable Aqua Dead Cell Stain Kit (Thermo Fisher Scientific) according to the manufacturer’s protocols. Cells were subsequently resuspended in staining buffer (PBS with 0.5% bovine serum albumin and 2 mM EDTA) and Fc-blocked using TruStain FcX reagent (BioLegend). Cells were then stained with the following fluorophore-conjugated anti-mouse antibodies for 30 min at 4°C: IL-15Rα-PE (clone DNT15Ra; Invitrogen), CD40-BV421 (clone 3/23; BD Biosciences), CD45-AF700 (clone 30-F11; Invitrogen), MHC Class II-APC-eF780 (clone M5/114.15.2; Invitrogen), CD11c-PE-eF610 (clone N418; Invitrogen), CD11b-PE/BV711/PE-Cy7 (clone M1/70; BioLegend, Invitrogen), XCR1-BV421/PE (clone ZET; BioLegend), SIRPα-BV605 (clone P84; BD Biosciences), Ly-6G-BV650/BV785 (clone 1A8; BioLegend), F4/80-AF488 (clone BM8; BioLegend), CD3e-PE-Cy5 (clone 145-2C11; Invitrogen), CD3-PE-Cy7 (clone 17A2; BioLegend), CD8a-SB780 (clone 53-6.7; Invitrogen), CD4-AF488 (clone GK1.5; Invitrogen), NK1.1-PE-Cy5/PE-eF610 (clone PK136; BioLegend, Invitrogen), CD19-PE-Cy5 (clone 1D3; Invitrogen), B220-PE-Cy5 (clone RA3-6B2; Invitrogen), CD44-BV650 (clone IM7; BioLegend), CD122-PerCP-Cy5.5 (clone TM-*β*1; BioLegend), CD27-APC (clone LG.3A10; BioLegend), KLRG1-BV711 (clone 2F1/KLRG1; BioLegend), CD62L-BV421 (clone MEL-14; BioLegend), and CXCR6-APC-Cy7 (clone SA051D1; BioLegend). Surface IL-15 was stained with biotinylated anti-mouse IL-15 (polyclonal rabbit; PeproTech) followed by streptavidin-APC (R&D). Data was acquired using an Attune NxT Flow Cytometer (Thermo Fisher Scientific) and analyzed using FCS Express flow cytometry analysis software (De Novo Software).

### Single cell RNA sequencing

ScRNA-seq was performed on freshly dissociated bulk tumor cells or on CD45+ and CD45 FACS-sorted freshly dissociated tumor cells using a Chromium controller (10X Genomics), as previously described. Briefly, gel beads in emulsion were generated, cells were lysed and barcoded cDNA amplified for 12 cycles. Amplified cDNA was fragmented and subjected to end-repair, poly A-tailing, adapter ligation, and 10x–specific sample indexing following the manufacturer’s protocol. Bioanalyzer (Agilent Technologies) and QuBit (ThermoFisher Scientific) were used to quantify the libraries, which were then sequenced in dedicated flowcells in paired-end mode on a HiSeq 2500 (Illumina) targeting a depth of 5×10^4^–1×10^5^reads per cell.

### Single-cell RNA sequencing analysis

Raw sequencing data were aligned and quantified using CellRanger against the provided GRCh38 human reference genome. Seurat was then used for all remaining steps. For each sample, cells were first selected as expressing less than 16–20% mitochondrial genes and displaying a minimum of 200–300 and a maximum of 2500–3500 features. Data were then log-normalized using a scale factor of 10,000. The 2,000 most variable features were then identified, data were scaled based on all the features, and principal component analysis was performed. Dimensionality of the dataset was then assessed using the JackStraw and ElbowPlot functions. Clusters were calculated and data dimensions were reduced using the t-SNE and UMAP methods.

### Statistical analysis

Data were analyzed using Prism software (GraphPad). Unpaired two-tailed *t* tests and ordinary one-way analysis of variance (ANOVA) with Tukey’s multiple comparisons tests were used to compare two groups and three or more groups, respectively. Unless otherwise indicated, error bars depict standard deviation. Kaplan-Meier method with log-rank tests were used for survival analysis. For all statistical tests, *P* values ≤ 0.05 were considered to be statistically significant, indicated as * *P* ≤ 0.05, ** *P* ≤ 0.01, *** *P* ≤ 0.001, and **** *P* ≤ 0.0001.

## Acknowledgments

The authors would like to thank Christopher Garris for invaluable assistance and discussions, as well as members of the J.V.R. laboratory for excellent technical assistance and helpful feedback, members of the Memorial Sloan Kettering (MSK) Genitourinary Oncology and Urology Services for thoughtful discussions, and Gil Redelman-Sidi and Michael Glickman for generously providing bladder cancer cell lines and technical advice. This work was supported, in part, by a Young Investigator Award from the American Society of Clinical Oncology/Conquer Cancer Foundation to J.L.W. (any opinions, findings, and conclusions expressed in this material are those of the authors and do not necessarily reflect those of the American Society of Clinical Oncology or Conquer Cancer), as well as the following grants from the National Institutes of Health (NIH): UL1TR001866 and KL2TR001865 from the National Center for Advancing Translational Sciences NIH Clinical and Translational Science Award Program to J.L.W. and D.A.K.; K08CA248966 to D.A.K.; F32CA250147 to C.S.G.; R01CA244327, R35CA196620, and P01CA190174 to J.V.R.; MSK Specialized Program of Research Excellence in Bladder Cancer P50CA221745 through a Developmental Research Program Award; and the National Cancer Institute Cancer Center Support Grant P30CA008748. The content is solely the responsibility of the authors and does not necessarily represent the official views of the NIH. Support was also provided by the Bladder Cancer Advocacy Network through a Bladder Cancer Research Innovation Award; the V Foundation for Cancer Research under grant ID no. T2017-002; the Robertson Therapeutic Development Fund at Rockefeller University; and the American Society of Hematology through a Research Training Award for Fellows to D.A.K. We dedicate this study to the memory of Thomas Waldmann whose vision and commitment to immunotherapy inspired these studies.

## Figures and Tables

**Fig. S1.**
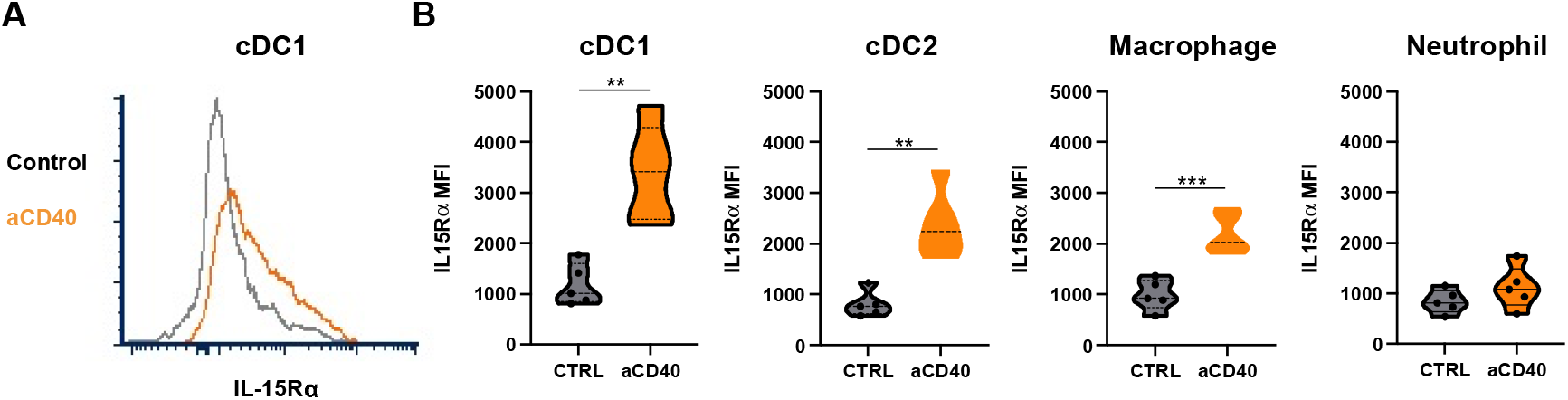
CD40 agonism induces IL-15Rα upregulation on dendritic cells in the UPPL1541 bladder tumor microenvironment. Mice bearing orthotopic UPPL1541 bladder tumors were treated intravesically with anti-CD40 antibody or isotype-matched control antibody on days 6 and 9 post-tumor implantation. (*A*) Representative histogram and (*B*) quantification across mice of IL-15Rα mean fluorescence intensity on type-1 conventional DCs (cDC1; defined as F4/80^−^Ly-6G^−^CD11c^+^MHCII^+^XCR1^+^), type-2 conventional DCs (cDC2; defined as F4/80^−^Ly-6G^−^CD11c^+^MHCII^+^SIRPα^+^), macrophages (defined as CD11b^+^F4/80^+^Ly-6G^−^), and neutrophils (defined as CD11b^+^Ly-6G^+^) in the bladder microenvironment as assessed by flow cytometry at day 12 post-tumor implantation (n = 5 mice per group). **p < 0.01, ***p < 0.001.

**Fig. S2.**
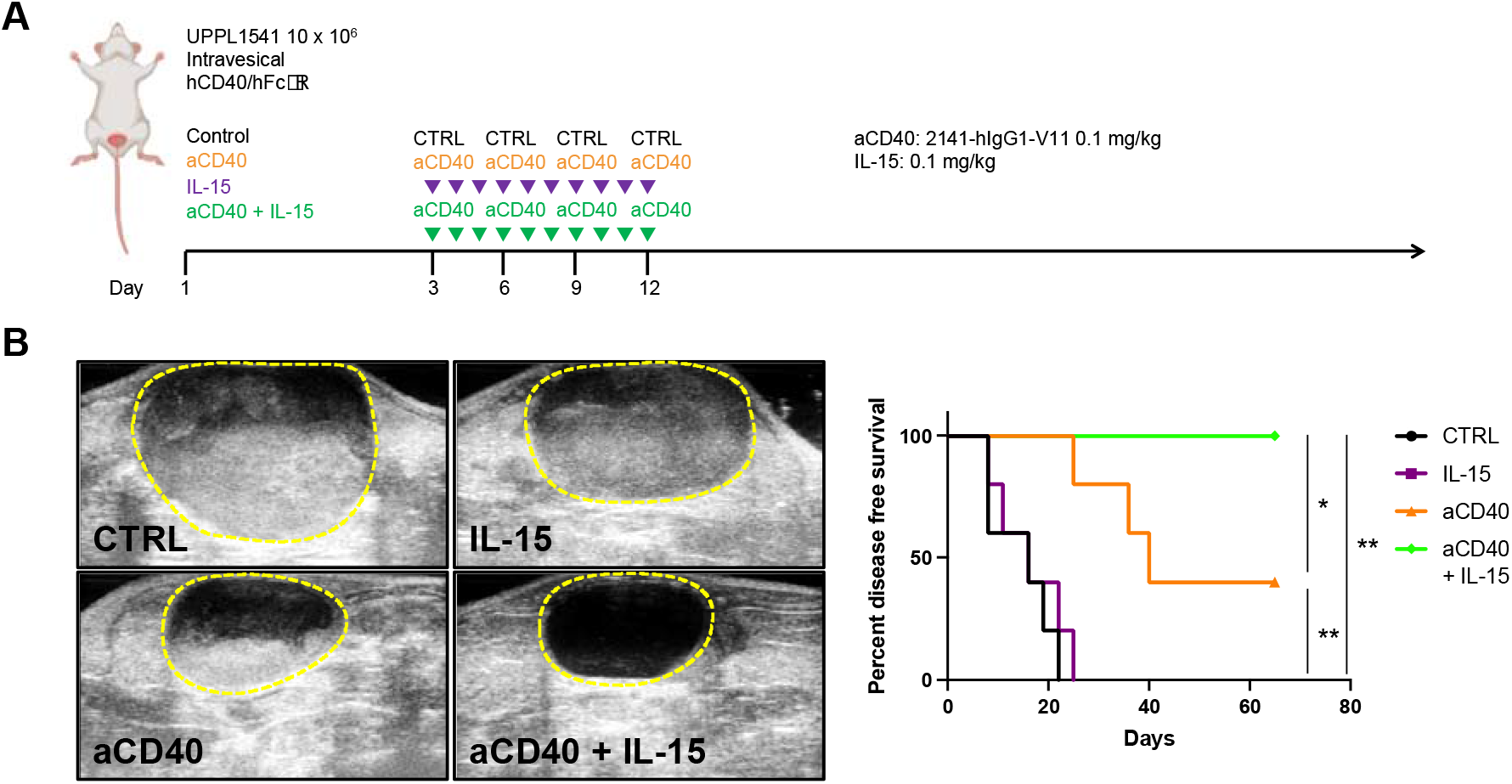
Combination of Fc-optimized anti-CD40 agonist antibody 2141-V11 and IL-15 enhances anti-tumor activity against UPPL1541 tumors. (*A*) Schematic of the treatment of humanized hCD40/hFcyR mice bearing orthotopic UPPL1541 bladder tumors with anti-CD40 antibody 2141-V11 and/or IL-15 or control (isotype-matched control antibody and/or vehicle). (*B*) Representative bladder ultrasound imaging (left) at day 45 post-tumor implantation and disease-free survival (right) of mice treated as outlined in A. *p < 0.05, **p < 0.01.

**Fig. S3.**
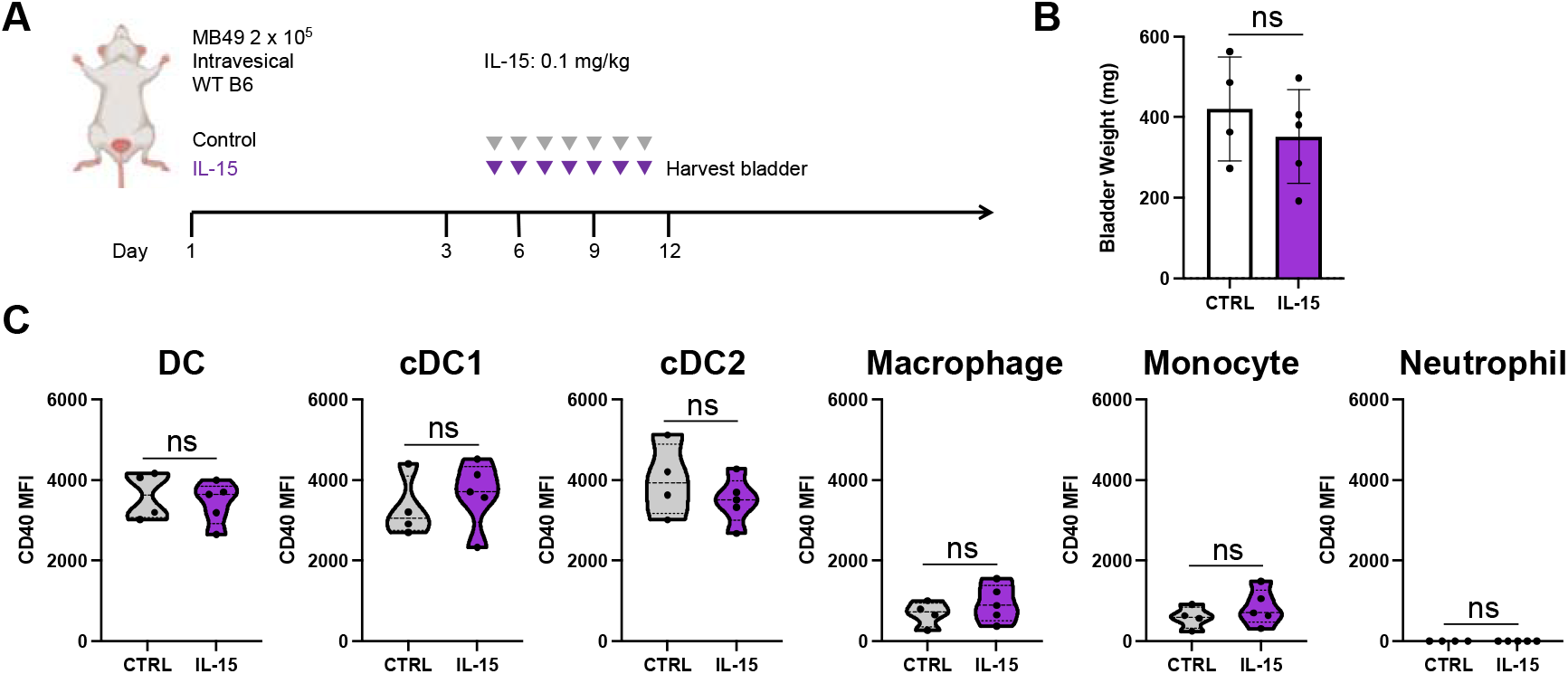
IL-15 therapy does not modulate myeloid CD40 expression. (*A*) Schematic of the treatment of mice bearing orthotopic MB49 bladder tumors with IL-15 or vehicle control. (*B*) Bladder weights across mice at day 12 post-tumor implantation (n = 4-5 mice per group; bars represent SD). (*C*) CD40 mean fluorescence intensity across mice on total dendritic cells (DC; defined as F4/80^−^Ly-6G^−^CD11c^+^MHCII^+^), type-1 conventional DCs (cDC1; defined as F4/80^−^Ly-6G^−^CD11c^+^MHCII^+^XCR1^+^), type-2 conventional DCs (cDC2; defined as F4/80^−^Ly-6G^−^CD11c^+^MHCII^+^SIRPα^+^), macrophages (defined as CD11b^+^F4/80^+^Ly-6G^−^), monocytes (defined as CD11b^hi^Ly6G^−^), and neutrophils (defined as CD11b^+^Ly-6G^+^) in the bladder microenvironment as assessed by flow cytometry at day 12 post-tumor implantation (n = 4-5 mice per group). ns = not significant.

## Notes

### Competing Interest Statement

The authors have declared no competing interest.

